# Sustained experimental activation of FGF8/ERK in the developing chicken spinal cord reproducibly models early events in ERK-mediated tumorigenesis

**DOI:** 10.1101/2021.06.10.447891

**Authors:** Axelle Wilmerding, Lauranne Bouteille, Nathalie Caruso, Ghislain Bidaut, Heather Etchevers, Yacine Graba, Marie-Claire Delfini

**Author notes:** Co-senior authors (Yacine Graba, Marie-Claire Delfini).

## Abstract

Most human cancers demonstrate activated MAPK/ERK pathway signaling as a key tumor initiation step, but the immediate steps of further oncogenic progression are poorly understood due to a lack of appropriate models. Spinal cord differentiation follows caudal elongation in vertebrate embryos; both processes are regulated by a FGF8 gradient highest in neuromesodermal progenitors (NMP), where kinase effectors ERK1/2 maintain an undifferentiated state. FGF8/ERK signal attenuation is necessary for NMPs to progress to differentiation. We show that sustained ERK1/2 activity, using a constitutively active form of the kinase MEK1 (MEK1ca) in the chicken embryo, reproducibly provokes neopasia in the developing spinal cord. Transcriptomic data show that neoplasia not only relies on the maintenance of NMP gene expression, and the inhibition of genes expressed in the differentiating spinal cord, but also on a profound change in the transcriptional signature of the spinal cord cells leading to a complete loss of cell-type identity. MEK1ca expression in the developing spinal cord of the chicken embryo is therefore a tractable *in vivo* model to identify the critical factors fostering malignancy in ERK-induced tumorigenesis.

## INTRODUCTION

The spinal cord is the most caudal part of the nervous system of vertebrates. The embryonic neural tube, from which it derives, is a pseudostratified epithelium generated progressively by neuromesodermal progenitors (NMP). These initially present mixed neural and mesodermal characteristics and are located in the most caudal part of the embryo (Henrique et al., 2015; Row et al., 2016; Wilson et al., 2009). Neurulation closely follows the caudal regression of the primitive streak and body elongation during the later stages of gastrulation. Several processes related to spinal cord specification and maturation are coupled to this caudal extension, including neurogenesis, ventral patterning, and neural crest specification; all these are regulated initially by Fibroblast Growth Factor 8 (FGF8) signaling, active in the NMP region and which maintains an undifferentiated state. Blocking FGF8 signaling in the caudal part of the embryo accelerates the onset of neural differentiation genes while ectopic maintenance of FGF8 inhibits neural differentiation (Bertrand et al., 2000; Diez del Corral et al., 2003; Patel et al., 2013). This gradient of FGF8 signaling also controls progressive maturation of the paraxial mesoderm, the other major derivative of the NMP, where overexpression of FGF8 in the presomitic mesoderm cells prevents its differentiation and segmentation (Dubrulle et al., 2001).

FGF signaling activates a variety of downstream effectors, including mitogen-activated protein kinase (MAPK) (extracellular signal-regulated kinase (ERK1/2)), and phosphatidylinositol 3-kinase (PI3K) (Böttcher and Niehrs, 2005). During early stages of chicken embryo development, ERK1/2 is the main cellular effector of FGF signaling. From the early blastula to the 10 somite stage, nearly all regions of ERK1/2 activity respond to FGF signaling (Lunn et al., 2007). We have shown that ERK1/2 is the effector of the gradient of FGF8 in the presomitic mesoderm that control the maturation of the paraxial mesoderm in chicken embryo (Delfini et al., 2005).

The transduction of many growth factor signals into a recipient cell normally occurs in the sequential activation, by phosphorylation, of RAS-like GTPases, RAF and MEK enzyme family members. This process converges on the interaction of activated ERK1/2 with hundreds of cytosolic and nuclear substrates to control key cellular events including cell cycle progression, proliferation, survival and differentiation (Samatar and Poulikakos, 2014). ERK1/2 activation plays an important role in the initiation of most cancers (Hoshino et al., 1999; Maik-Rachline and Seger, 2016). For this reason, significant efforts have been made to develop inhibitors of this pathway, several of which are used in chemotherapy (Samatar and Poulikakos, 2014). However, while such MAPK pathway inhibitors have improved clinical outcomes overall, emergence of drug resistance often limits their therapeutic efficacy in time (Maik-Rachline and Seger, 2016). It remains necessary to identify new targets and molecular mechanisms of oncogenic ERK1/2 activity in order to open further therapeutic perspectives.

In this study, we have used the chicken embryo to study the consequences of constitutive ERK1/2 activation in the developing spinal cord, by expressing a constitutively active form of MEK1 (MEK1ca), the kinase upstream of ERK1/2 (Delfini et al., 2005; Mansour et al., 1994) and comparing its effects to those of sustained FGF8. We have confirmed that the ERK1/2 pathway is the effector of FGF8 signaling in the control of neural differentiation in the developing spinal cord. Ectopic ERK1/2 activation leads to cell-autonomous re-expression of typical NMP genes and repression of usual markers of the differentiating spinal cord. Furthermore, MEK1ca expression in the developing spinal cord rapidly and reproducibly induces neoplasia.

Since the neural tube of the trunk gives rise to both spinal cord (neurons and glia) and neural crest cells (from which melanocytes and sympathetic ganglia in particular are derived) (Le Douarin and Kalcheim, 1999), MEK1ca expression by electroporation into the chicken embryo neural tube is an easily accessible *in vivo* model of the molecular and cellular effects of oncogenic activation of MAPK/ERK signaling, relevant to a range of pediatric cancers including neuroblastoma, astrocytoma or glioblastoma, and to melanoma.

## RESULTS

### ERK1/2 mediates FGF8 signaling to maintain undifferentiated cell states in the developing spinal cord

Inhibiting ERK1/2 activity using the MEK1/2 inhibitor PD184352 completely downregulates the expression of FGF8 target genes such as SPRY2 in the caudal part of the chicken embryo, showing that ERK1/2 are the main effectors of FGF8 in this tissue (Martínez-Morales et al., 2011). To test if maintaining ERK1/2 activity phenocopies FGF8 gain-of-function in the control of neural differentiation onset in the developing spinal cord, we electroporated the trunk neural tube of 2-day-old chicken embryos *in ovo* (Figure 1A) with a vector expressing a constitutively active form of MEK1, a kinase encoded by the gene *MAP2K1*, directly responsible for the phosphorylation and activation of ERK1/2 (MEK1ca (Delfini et al., 2005; Mansour et al., 1994)) or with a vector expressing FGF8 (Delfini et al., 2009). These vectors co-express GFP, and the control pCIG vector expresses only a nuclear-targeted form of GFP. Immunofluorescence on transverse sections with the phospho-Thr202/Tyr204-ERK1/2 (pERK) antibody one day after electroporation show that both FGF8 and MEK1ca expression in the trunk neural tube leads to a strong increase in activated ERK1/2 signaling (Figure 1B, Figure S1). As expected, this activation appeared respectively indirect (non-cell-autonomous, for FGF8) or direct (cell-autonomous, for MEK1ca). Using whole mount and transverse section *in situ* hybridization and immunostaining at this time, we observe that one day after electroporation, genes expressed in the NMP and in the most posterior part of the neural tube (such as SPRY2, CDX4 and GREB1), where ERK1/2 is activated in wild type embryo (Lunn et al., 2007), are upregulated after FGF8 and MEK1ca expression in the trunk neural tube. Conversely, genes expressed in the most anterior part of the trunk neural tube (such as PAX6, NKX6.2 and WNT4) are downregulated by ectopic FGF8 and MEK1ca expression (Figure 1, Figures S1-6). The effect on these target transcripts was likewise long range and non-cell autonomous for the FGF8 ligand, but cell-autonomous for MEK1ca. These results are consistent with a physiological function for MEK1 and ERK1/2, downstream of FGF signaling during neural tube development, in maintaining cells of the most caudal part of the embryo in a progenitor state.

**Figure 1:**
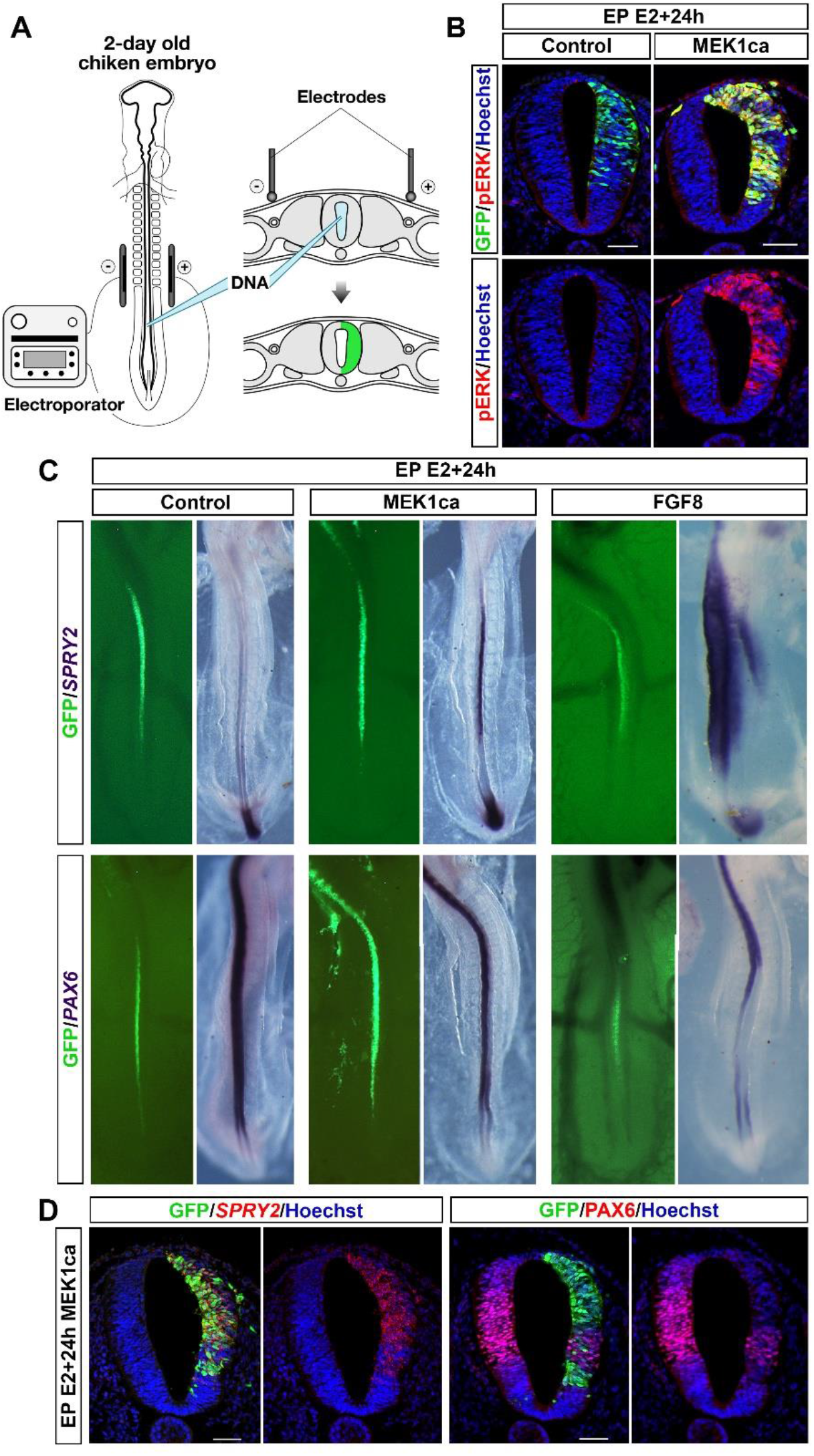
MEK1ca expression phenocopies FGF8 gain-of-function in a cell-autonomous manner. **A-** The transfection of the neural tube of a two-day-old chicken embryo is done by unilateral or bilateral electroporation with vectors expressing GFP (control) or co-expressing GFP with MEK1ca or FGF8. **B-** Immunofluorescences on transverse sections with anti-pERK and anti-GFP antibodies one day post-electroporation with the control (pCIG) or MEK1ca-expressing vectors. **C-** Dorsal view of electroporated embryos one day post-electroporation with the control (pCIG), MEK1ca-, or FGF8-expressing vectors. Whole mount *in situ* hybridizations with SPRY2 and PAX6 probes, with corresponding GFP expression on the left. **D-** Fluorescent *in situ* hybridization with SPRY2 probe and immunofluorescence using anti-GFP and anti-PAX6 antibodies on trunk-level transverse sections of a chicken embryo one day postelectroporation with the MEK1ca-expressing vector. Blue is Hoechst staining of nuclear DNA. Scale bar: 50μm.

### MEK1ca expression in the chicken neural tube leads to neoplasia

In addition to the expected molecular phenotype obtained after MEK1ca transfection, which mimics FGF8 gain-of-function (Bertrand et al., 2000; Diez del Corral et al., 2003), we found that expression of MEK1ca leads to a profound neuroepithelial disorganization. Morphological changes in the neural tube can be observed as early as one day post-electroporation (Figure 2 and Figure S7-8). The phenotype becomes striking by the second day after transfection, with MEK1ca-expressing cells forming disorganized aggregate structures evoking solid tumors, both at the transfected site, with a greater effect on the dorsal region of the neural tube, and also at a distance from the electroporation site, particularly in the epidermis, the head, the heart, and also in the extraembryonic annexes, evoking metastases (Figure 2A-C and Figure S7, no distant GFP+ aggregates were observed in ten embryos transfected by pCIG, whereas eleven distant GFP+ aggregates were found in six MEK1ca embryos in a blinded experiment (0/10 for pCIG versus 11/6 for MEK1ca)). F-ACTIN visualization with fluorescence-conjugated phalloidin, and immunofluorescence for a mitotic marker, pS28-H3 (pHH3), together highlight the disorganization of the neural tube on the electroporated side of MEK1ca-transfected embryos (Figure 2D-E, Figure S8A). Indeed, mitotic cells are not restrained to the apical part of the future neural ependyma, as is the case in the control condition. Immunostaining with antibodies against the progenitor and neuronal differentiation markers SOX2 and TUJ1, respectively, and *in situ* hybridization with a pan-neuronal *SCG10* probe highlights that MEK1ca inhibits differentiation in the neural tube (Figure 2F-G and Figure S8B). Cleaved caspase-3 (CASP3) immunofluorescence shows that MEK1ca expression also leads to an increase of cell death in the electroporated side after 24h and in some of the tumor-like structures at later time points (Figure S9). The increase of apoptosis in the tumors is consistent with previous studies showing that ERK activity, in addition to its well-described oncogenic and hyperproliferative effects, also promotes apoptosis, autophagy and senescence (Cagnol and Chambard, 2010). MEK1ca also results in cells invading the neural tube lumen (Figure 2E’). Multinucleated cells could be observed in some cases (Figure 2H), suggesting that induction of multinucleation by RAS/ERK overactivation (Dikovskaya et al., 2015) may be an early event during ERK-induced tumorigenesis.

**Figure 2:**
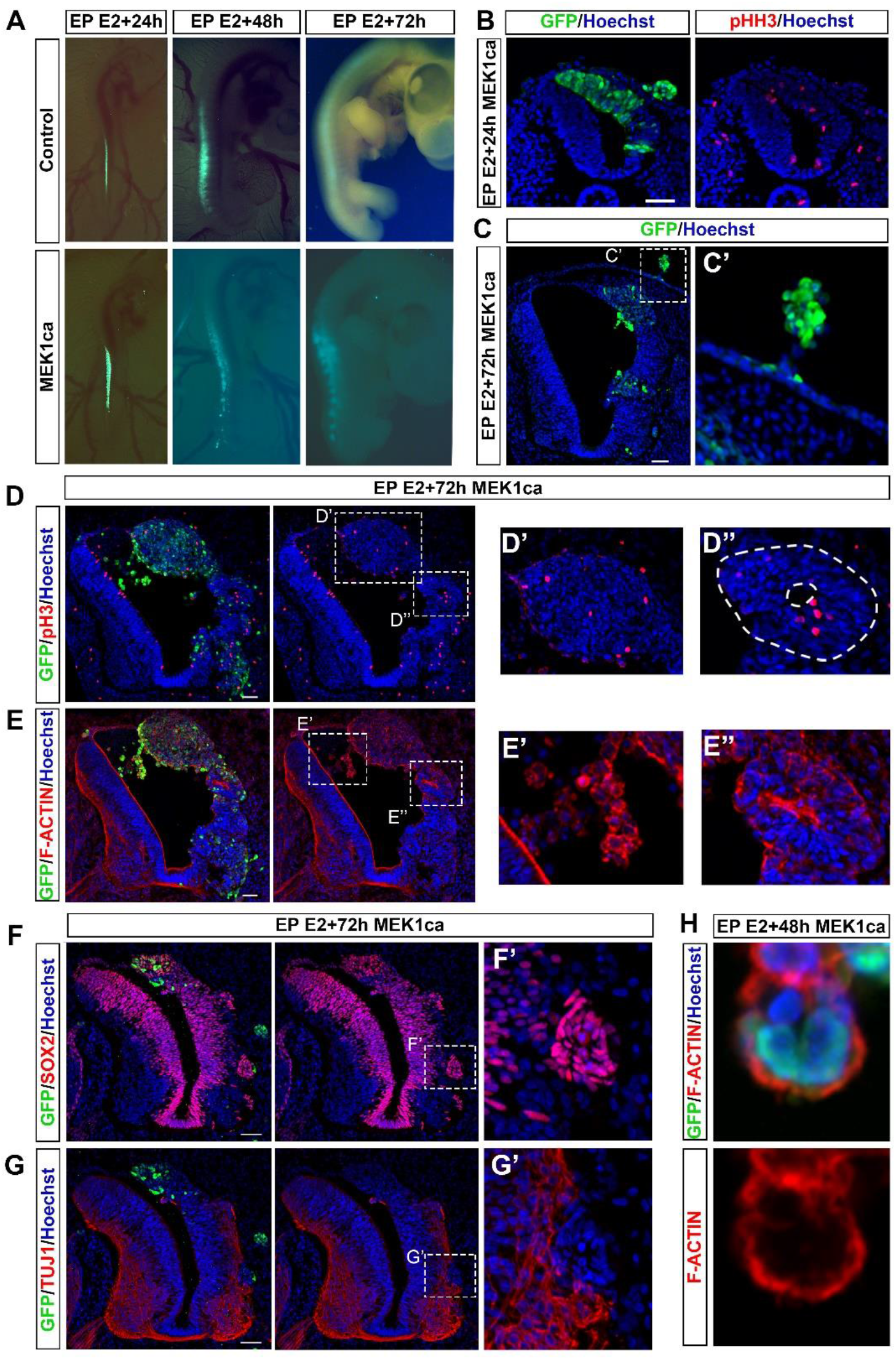
MEK1ca expression in the differentiating neural tube leads to neoplasia. **A-** View of whole chicken embryo showing GFP fluorescence at one, two, or three days postelectroporation with control (pCIG) or MEK1ca-expressing vectors. **B-** Immunofluorescence on transverse sections at the trunk level with anti-GFP and anti-pHH3 antibodies (metaphase marker) one day post-electroporation with the MEK1ca-expressing vector. **C-** Immunofluorescence on transverse sections with anti-GFP, three days post-electroporation with the MEK1ca-expressing vector. **C’** is higher magnification within **C** highlighting an abnormal growth in the epidermis formed by GFP-positive cells. **D-G’-** Immunofluorescence on transverse sections at 3 days post-electroporation with the MEK1ca-expressing vector, using antibodies against GFP in conjunction with pHH3 (**D**), SOX2 (a neural progenitor marker) (**F**) and TUJ1 (a neuronal differentiation marker) (**G**) or F-ACTIN phalloidin staining (**E**). **D’** and **D’’** highlight ectopic mitosis both in the most dorsal part of the neural tube (**D’**) and in the rest of the tube (**D’’**), forming rosette-like structures highlighted by F-ACTIN staining (**E’’**). **E’** shows transfected cells invading the neural tube lumen. **F’** and **G’** focus on an aggregate of GFP-positive cells in the differentiated area of the neural tube which retain SOX2 immunoreactivity and are negative for TUJ1. **H-** Immunofluorescence on a transverse section with anti-GFP and F-ACTIN labeling, two days post-electroporation with the MEK1ca-expressing vector, showing a cell in the lumen of the neural tube with multiple nuclei. Blue is Hoechst staining of nuclear DNA. Scale bar: 50 μm.

### Transcriptomic data matches with the developmental function of ERK1/2 in the developing spinal cord

To investigate further the mechanisms underlying MEK1ca-induced neoplasia, we aimed at identifying the global transcriptional response induced by MEK1ca in the chicken neural tube. We performed an RNA-seq on GFP positive cells of the chicken embryo neural tube after MEK1ca electroporation (Figure 3A-B and Figure S10). E2 neural tubes were bilaterally electroporated with either the control vector pCIG or the MEK1ca expression vector (Figure 3A). The regions of the neural tube expressing the GFP were dissected one day after electroporation and dissociated. GFP-expressing cells were isolated by FACS. Two independent RNAs samples were extracted, reverse transcribed, and cDNAs were amplified using linear amplification and used for sequencing library building. RNA-seq data from alignment to the Galgal4 genome assembly identified, for FDR5 (False Discovery Rate = 5%), 2316 genes with significantly changed expression (Figure 3B, Supplementary Table 1 and Supplementary Table 2; see also Material and Methods section), of which 1310 were up-regulated (57%) (Supplementary Table 1) and 1006 down-regulated (43%) (Supplementary Table 2) (Figure 3C top panel). The tendency to act as activator rather than repressor is increased when selecting genes differentially expressed by more than two-fold, with 348 being up-regulated (86 %) and only 57 down-regulated (14 %) (Figure 3C bottom panel). We validated the transcriptomic data by *in situ* hybridization with probes for 5 upregulated genes (Supplementary Table 5): CLDN1 (Figure 3E), IL17RD, EGR1, LIN28A and CHST15 probes (Figure S11). The transcriptomic phenotype is homogeneous throughout the neural tube, with all MEK1ca positive cells behaving similarly, including in the dorsal most part of the neural tube containing neural crest cells (Figure 3E and Figure S11).

Panther overrepresentation analysis for biological process (Mi et al., 2013) of the transcriptomic data for all the upregulated or all the downregulated genes (FDR = 5%) (Figure 3F,G and Supplementary Table 3 and 4) matches with the well-known autoregulatory loop of the MAPK/ERK pathway (Kidger and Keyse, 2016), the cross talk of the MAPK/ERK pathway with the PI3K pathway (Won et al., 2012), and with classical oncogenic functions of MAPK/ERK pathway such as the control of extra-cellular matrix and angiogenesis (Dias Carvalho et al., 2018; Reddy, 2003). RNA-seq results also matches the developmental function of ERK1/2 downstream of FGF8 in controlling the onset of neural differentiation in the developing spinal cord (Figure 4A): genes known to be endogenously expressed in the most caudal part of the embryo are upregulated (i.e. SPRY2, CDX4, GREB1, DUSP4, DUSP5, DUSP6, IL17RD, CLDN1 and LIN28A genes) and genes endogenously expressed in the differentiating neural tube are downregulated (i.e. PAX6, DBX2, WNT4, ENPP2, NKX6.2 and PLXNA2 genes) (Olivera-Martinez et al., 2014) (Figure 4B-C, Supplementary Tables 1-2).

**Figure 3:**
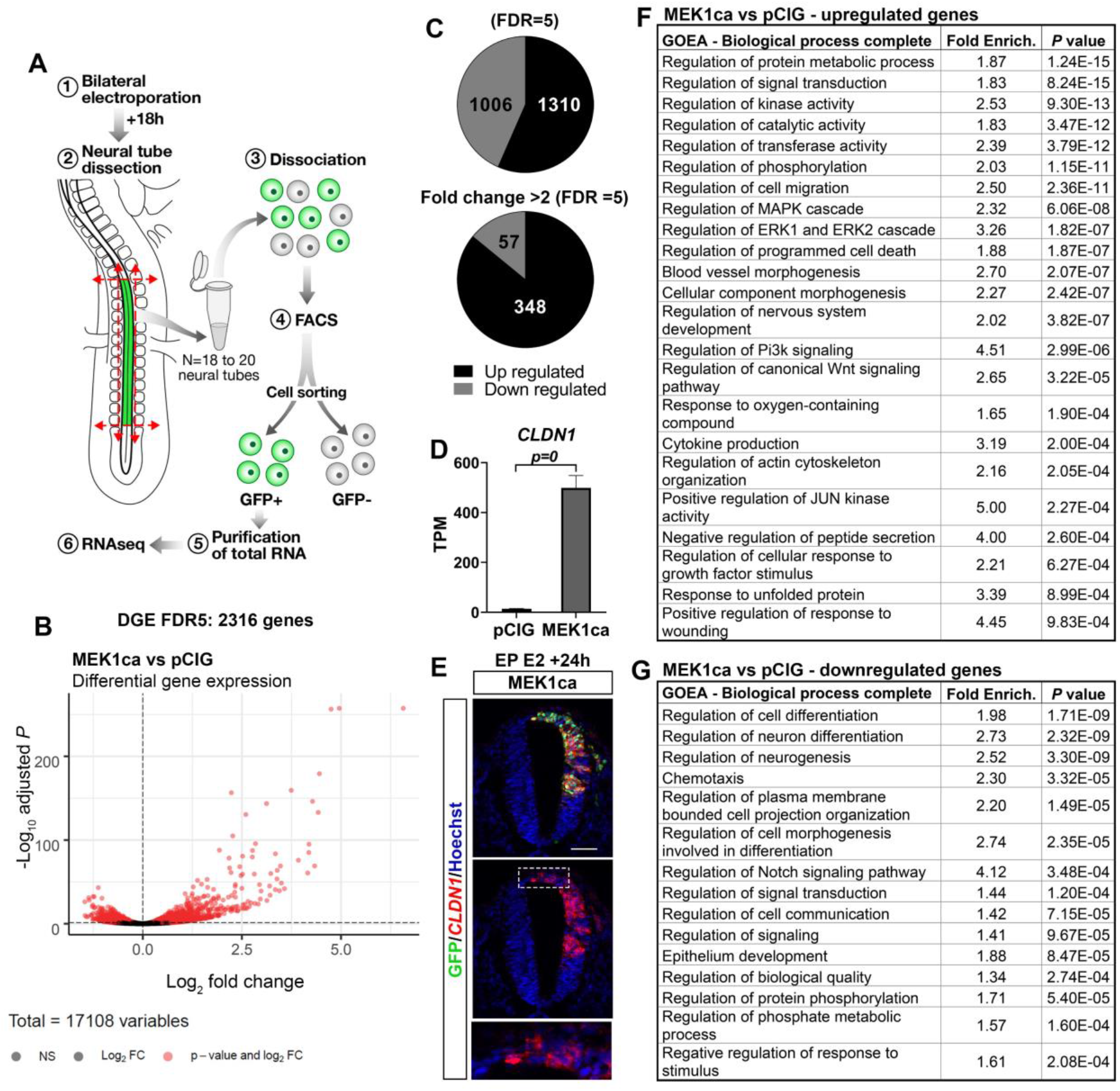
Characterization of the MEK1ca-induced transcriptome. **A-** One day after bilateral electroporation of the trunk neural tube at Hamburger-Hamilton (HH) stage HH12 with the pCIG control or the MEK1ca vector, the electroporated region of the neural tube was dissected (18-20 embryos per condition in duplicate) and GFP-positive cells were sorted by FACS. **B-** Volcano plot of differential gene expression (DGE) for the MEK1ca versus control conditions. **C-** Pie charts representing the number of upregulated and downregulated genes in the presence of MEK1ca for FDR=5% (all the genes) or for FDR=5% and with greater than twofold change in normalized transcript numbers. **D-** Graph of *CLDN1* TPM (transcripts per kilobase million), obtained for the two replicates of the control (pCIG1, pCIG2) and MEK1ca-(MEK1ca-1 and MEK1ca-2) expressing samples. **E-** Fluorescent *in situ* hybridization with *CLDN1* probe and immunofluorescence with anti-GFP antibody on trunk-level transverse sections of a chicken embryo one day post-electroporation with the MEK1ca-expressing vector. Blue is Hoechst staining of nuclear DNA. Scale bar: 50 μm. **F-G-** Gene Ontology enrichment analysis (GOEA) of the biological processes for up- (**F**) and downregulated (**G**) genes in the MEK1ca versus control conditions (FDR=5%).

### MEK1ca expression in the neural tube induces cell-type inappropriate transcription

In addition to being consistent with the normal developmental role of ERK1/2 during the spinal cord differentiation process, transcriptomic data highlight that MEK1ca leads to the ectopic expression of genes that are neither expressed in NMP nor in the neural tube at that stage (e.g. IL1R1, IL1RL2, B3GNT, RGS8, AQP1, ARAP3 and FAM65B genes). *In situ* hybridization to *AQP1* confirmed the transcriptomic data (Figure 4E), showing ectopic expression triggered by MEK1ca in cells of the trunk neural tube, including in the most dorsal part (Figure 4F). Therefore, neoplasia induced by MEK1ca expression might not only be due to the maintenance of an immature NMP cell state, but also implies aberrant cell fating.

**Figure 4:**
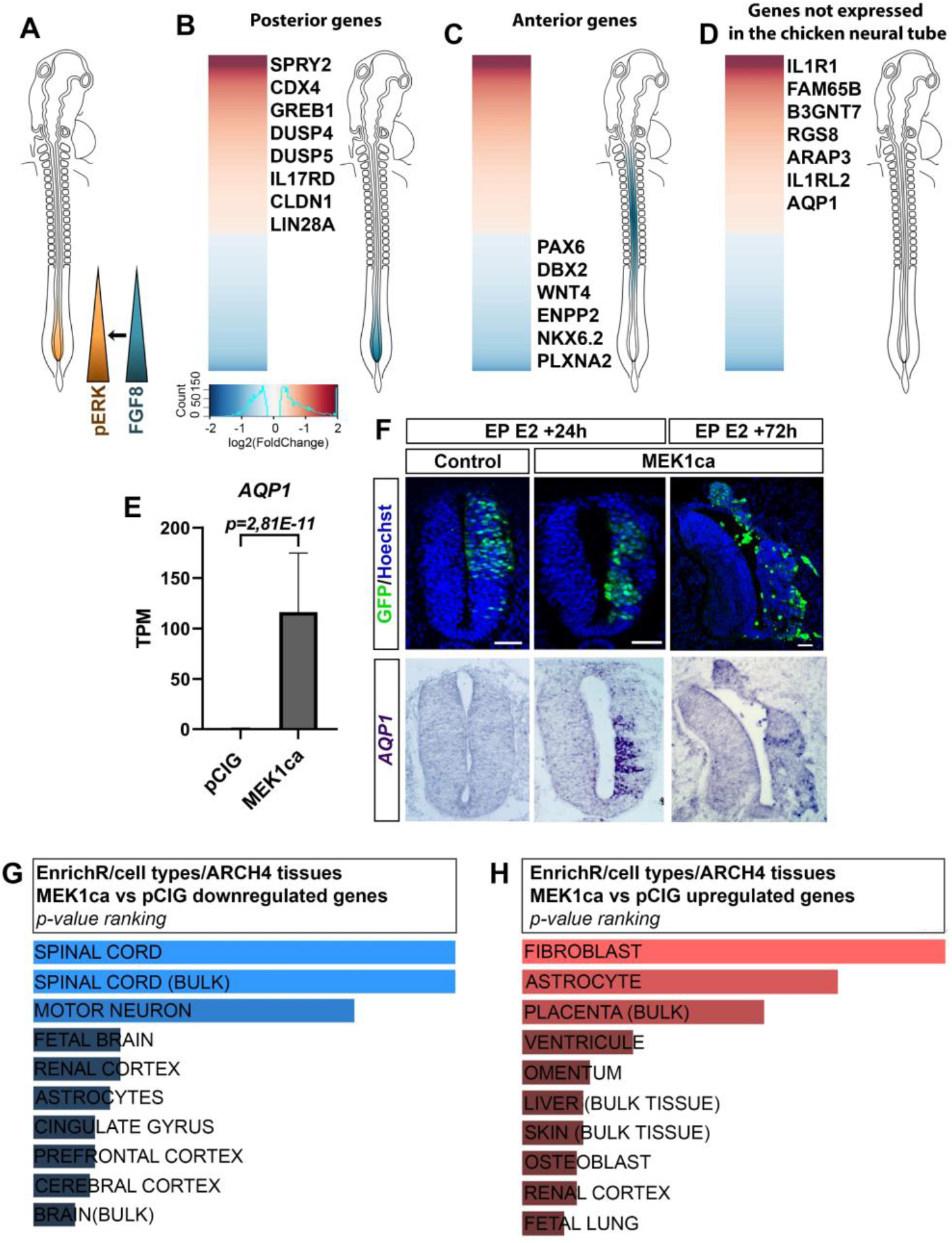
Expected and unexpected features of the MEK1ca-induced transcriptome. **A-** Diagram showing the expression and activation domains of FGF8 and ERK1/2 respectively in the trunk of chicken embryo at E2. **B-D-** Heat map of 2316 genes deregulated in MEK1ca-versus pCIG-transfected embryos (DGE with FDR5=5%) with a list of typical (**B**) posterior genes that are upregulated, (**C**) anterior genes that are downregulated, and (**D**) genes not usually expressed in the trunk of chicken embryo that are ectopically upregulated. **E-** Graph of the mean *AQP1* TPM (transcripts per kilobase million), obtained for the replicates of the control (pCIG1, pCIG2) and MEK1ca-(MEK1ca-1 and MEK1ca-2) expressing samples. **F-** *In situ* hybridization with *AQP1* probe and immunofluorescence using an anti-GFP antibody on trunk-level transverse sections of chicken embryos one or three days post-electroporation with the MEK1ca-expressing vector. Blue in top panel is Hoechst staining of nuclear DNA. Scale bar: 50 μm. **G-H-** Comparison of the lists of downregulated (**G**) and upregulated (**H, I**) genes for MEK1ca-vs pCIG-transfected neural tubes with the ARCHS4 Tissues database (**G-H**) and the EnrichR analysis tool for cell types.

Comparing the list of MEK1ca-deregulated genes with the ARCHS4 tissue database (Lachmann et al., 2018) using the “EnrichR” analytical tool (Chen et al., 2013; Kuleshov et al., 2016; Xie et al., 2021) showed that as expected, genes downregulated correlate with spinal cord identity (Figure 4G, Figure S12A). Upregulated genes found in this comparison are instead associated with fibroblast, placenta, liver, osteoblast, and renal tissues (Figure 4H, Figure S12B). We interpret these results as indicative of aberrant cell fates, a feature of cancer cells (Suzuki et al., 2014), which our study further suggests may occur very early in tumorigenesis following ERK hyperactivation.

The *in ovo* model we describe here develops tumor-like structures as early as the next day. By two days post-transfection, tumors are found not only in the neuroepithelium, at the site of the transfection, but also in adjacent tissues like the epidermis and as distant as the extraembryonic annexes. MEK1ca takes two weeks to induce palpable tumors in nude mice (Mansour et al., 1994). The speed of tumor appearance and the widespread dissemination of transfected cells throughout the technically accessible chicken embryo are experimentally advantageous.

In conclusion, MEK1ca electroporation into the trunk neural tube of a two-day-old chicken embryo is an attractive *in vivo* vertebrate model of ERK-mediated oncogenic activation that is rapid, cost-effective, and allows study from the onset of both the cell-autonomous and non-cell-autonomous effects of ERK overactivation. Furthermore, this model respects the 3D rule (Sneddon et al., 2017) in that at the stages analyzed, it is not subject to the same ethical constraints on animal experimentation as mouse models. While this chicken embryo model does not recapitulate a specific human cancer type, it addresses some general features of ERK-induced tumors. It might in particular be suited for studying aspects of pediatric central nervous system tumors (Kram et al., 2018), as well as features of two frequent cancers with neural crest origins in adults and children respectively, *i.e*. melanoma (Shakhova, 2014) and neuroblastoma (Tsubota and Kadomatsu, 2018). Our model is particularly suited to perform experiments in epistasis to understand how candidate proteins may interact with ERK-mediated MAPK pathways during tumorigenesis and to test the efficacy of pathway-inhibitory therapies.

## MATERIALS AND METHODS

### Ethics statement

Experiments performed with non-hatched avian embryos in the first twothird of embryonic developmental time are not considered animal experiments according to the directive 2010/63/EU.

### Chicken embryos

Fertilized chicken eggs were obtained from EARL les Bruyeres (Dangers, France) and incubated horizontally at 38°C in a humidified incubator. Embryos were staged according to the developmental table of Hamburger and Hamilton (HH) (Hamburger and Hamilton, 1992) or according to days of incubation (E).

### In ovo electroporation and plasmids

Neural tube *in ovo* electroporations were performed around HH11/12 as in (Delfini and Duprez, 2004). Eggs were windowed, and the DNA solution was injected in neural tube lumen. Needle L-shape platinum electrodes (CUY613P5) were placed on both sides of the embryo at trunk level (5 mm apart), with the cathode always at its right. Five 50 ms pulses of 25 volts were given unilaterally (or bilaterally for RNAseq experiments) at 50 ms intervals with an electroporator NEPA21 (Nepagene).

The plasmids used for the gain- and loss-of-function experiments co-express a cytoplasmic or nuclear GFP (pCAGGS and pCIG respectively, used alone as controls) and the coding sequence (CDS) of the gene of interest. Vector used were: pCIG-MEK1ca (Delfini et al., 2005) and pCAGGS-FGF8 (Delfini et al., 2009). The plasmids used for electroporation were purified using the Nucleobond Xtra Midi kit (Macherey-Nagel). Final concentration of DNA delivered per embryo for electroporation is between 1 to 2μg/μl.

### Immunofluorescence and fluorescent in situ hybridization

Embryos were fixed in 4% buffered formaldehyde in PBS then treated with a sucrose gradient (15% and 30% in PBS), embedded in OCT (optimal cutting temperature) medium and stored at −80°C. Embryos were sectioned into 16 μm slices with a Leica cryostat and the slides were conserved at − 80°C or directly used for FISH and/or immunofluorescence.

#### Immunofluorescence

Slides were rehydrated in PBS then blocked with 10% goat serum, 3% BSA, 0,4% Triton X100 in PBS for one hour. Primary antibodies were incubated overnight diluted in the same solution at 4°C. The following primary antibodies were used in this study: chicken anti-GFP 1:1000 (1020 AVES), rabbit anti-phospho-p44/42 MAPK (ERK1/2) (Thr202/Tyr204) 1:250 (Cell signaling #9101), rabbit anti-SOX2 1:500 (AB5603 Merck Millipore), mouse anti-TUJ1 1:500 (801202 Biolegend), rat anti-pHH3 1: 250 (S28, Abcam ab10543), rabbit anti-cleaved-caspase 3 1:500 (Asp175, CST 9661), mouse anti-PAX6 1:500 (Developmental Studies Hybridoma Bank), Phalloidin AlexaFluor 568 1:40 (Thermofisher). The secondary antibodies used were: anti-chicken, anti-rabbit, anti-mouse or anti-rat with conjugated fluorochromes (Alexa Fluor 488, 568 or 647; ThermoFisher) at 1:500. Sections were incubated for one hour in the blocking solution containing Hoechst dye (1:1000). Slides were washed, mounted (Thermo Scientific Shandon Immu-Mount) and imaged with a Zeiss microscope Z1 Apotome or a confocal LSM 780.

#### Fluorescent in situ hybridization on tissue section

The protocol for fluorescent *in situ* hybridization on tissue section is as described (Delfini et al., 2013). Briefly, slides were treated with proteinase K 10 μg/ml (3 minutes at 37°C) in a solution of Tris-HCl 50 mM pH 7.5, then in triethanolamine 0.1M and 0.25% acetic anhydride. They were pre-incubated with hybridization buffer (50% formamide, SSC 5X, Denhardt 5X, yeast tRNA 250 μg/ml and herring sperm DNA 500 μg/ml) for 3 hours at room temperature, then incubated in the same buffer with digoxygenin (DIG)-labelled RNA probes overnight at 55°C in a humid chamber. The slides were then washed twice with 0.2X SSC for 30 minutes at 65°C. After 5 minutes in TNT buffer (100 mM Tris pH7.5, 150mM NaCl and 0.1% Tween-20), they were blocked for 1 hour in buffer containing 1X TNT, 1% Blocking Reagent (BR, Merck 11096176) and 10% goat serum, then incubated in the same buffer for 3h with anti-DIG-peroxidase antibodies (1:500, Roche) and revealed using the TSA-Plus Cyanine-3 kit (Perkin-Elmer). RNA probes used for *in situ* hybridization were: SPRY2, PAX6, NKX6.2, WNT4, GREB1, CDX4, SCG10, CLDN1, and AQP1 (primers used to generate the probes are listed in Table 5). The plasmids used to generate the CLDN1 and *SCG10* RNA probes were kind gifts from Jean-Loup Duband and Hermann Rohrer, respectively. All the other probes were produced from PCR products amplified from cDNA of E3 chicken embryo neural tubes (either WT or transfected by MEK1ca). PCR primer pairs used for each probe are in Supplementary Table 3.

### Whole mount in situ hybridization

The whole mount *in situ* hybridization protocol is as described (Wilmerding et al., 2021). Embryos were fixed 2 hours at RT in 4% buffered formaldehyde in PBS. Embryos were dehydrated with sequential washes in 50% ethanol/ PBS+ 0.1% Tween20 then 100% ethanol and conserved at −20°C. Embryos were bleached for 45 minutes in 80% ethanol, 6% aqueous H2O2 and then rehydrated. They were treated 10 minutes with proteinase K 10 μg/ml at RT and refixed with 4% formaldehyde, 0.2% glutaraldehyde. After 1 hour of blocking in the hybridization buffer (50% formamide, SSC 5x, 50 μg/mL heparin, yeast tRNA 50 μg/mL, SDS 1%), hybridization with DIG-labelled RNA probes was performed at 68°C overnight. The next day, embryos were washed in hybridization buffer then once in TBS (25 mM Tris, 150 mM NaCl, 2 mM KCl, pH 7.4) +0.1% Tween 20. They were incubated 1h at RT in a blocking buffer (20% BR + 20% goat serum) and then overnight with an anti-DIG-AP antibody (1:2000, Merck) in the blocking buffer. After 3 washes (1 hour) in TBS+0.1% Tween 20, embryos were equilibrated in NTMT buffer (NaCl 100 mM, TrisHCl 100mM pH 9.5, MgCl2 50 mM, 0.2% Tween20) and incubated in NBT/BCIP (Promega) at RT in the dark until color development. Pictures of whole embryos were taken with a BinoFluo MZFLIII and a color camera.

### RNA-seq analysis

Electroporations were carried out as described above with 5 bilateral pulses. Plasmid DNA concentrations were, for the control mix, pCIG at 2 μg/μl, and for the MEK1ca mix at pCIG-MEK1ca 1 μg/μl + pCIG 1μg/μl. Parts of the neural tube expressing GFP were microdissected one day after electroporation and dissociated with trypsin/EDTA 0.25%. A highly enriched population of GFP-expressing cells was isolated by FACS with the use of a dead cell exclusion (DCE)/discrimination dye (DAPI) to eliminate dying cells (Wilmerding et al., 2021). RNA was extracted (RNeasy Mini Kit), reverse transcribed with random primers and cDNA was amplified using a linear amplification system used for building sequencing libraries (done at the GATC). After adapter ligation and adapter-specific PCR amplification, libraries were sequenced on an Illumina NextSeq for a total of 50,000,000 paired end reads with 2 x 50 bp read length. Bioinformatics analyses were done using the galgal4.0 chicken genome. Qualitative analysis of RNA-seq data from the two biological replicates shows a high Pearson correlation score (>0,99), indicating experimental reproducibility (Figure S10).

### Quantifications and statistical significance

All in situ hybridization experiments and immunofluorescence stains at each stage were done on a minimum of three embryos (biological replicates).

## Acknowledgements

We sincerely thank Dr Andrew Saurin for his help in the RNA-seq analysis. We would like to thank Sophie Gournet (IBPS, CNRS UMR7622) for the drawings in Figures 1, 3 and 4. We thank Dr Jean-Loup Duband and Dr Hermann Rohrer for their generous gifts of RNA probes. We sincerely thank Dr Samuel Tozer for his critical reading of the manuscript. We thank the Optical Imaging Platform of the IBDM. FACS experiments were undertaken with assistance from the CRCM (Marseille, France). RNA-seq sequencing were undertaken by the GATC. We are grateful to the Canceropôle PACA, the ANR, the IBDM and La Ligue contre le Cancer for their funding support.

## REFERENCES

Bertrand, N., Médevielle, F., and Pituello, F. (2000). FGF signalling controls the timing of Pax6 activation in the neural tube. Dev. Camb. Engl. 127, 4837–4843.

Böttcher, R.T., and Niehrs, C. (2005). Fibroblast Growth Factor Signaling during Early Vertebrate Development. Endocr. Rev. 26, 63–77.

Cagnol, S., and Chambard, J.-C. (2010). ERK and cell death: Mechanisms of ERK-induced cell death - apoptosis, autophagy and senescence: ERK and cell death. FEBS J. 277, 2–21.

Chen, E.Y., Tan, C.M., Kou, Y., Duan, Q., Wang, Z., Meirelles, G.V., Clark, N.R., and Ma’ayan, A. (2013). Enrichr: interactive and collaborative HTML5 gene list enrichment analysis tool. BMC Bioinformatics 14, 128.

Delfini, M.-C., and Duprez, D. (2004). Ectopic Myf5 or MyoD prevents the neuronal differentiation program in addition to inducing skeletal muscle differentiation, in the chick neural tube. Development 131, 713–723.

Delfini, M.-C., Dubrulle, J., Malapert, P., Chal, J., and Pourquie, O. (2005). Control of the segmentation process by graded MAPK/ERK activation in the chick embryo. Proc. Natl. Acad. Sci. 102, 11343–11348.

Delfini, M.-C., De La Celle, M., Gros, J., Serralbo, O., Marics, I., Seux, M., Scaal, M., and Marcelle, C. (2009). The timing of emergence of muscle progenitors is controlled by an FGF/ERK/SNAIL1 pathway. Dev. Biol. 333, 229–237.

Delfini, M.-C., Mantilleri, A., Gaillard, S., Hao, J., Reynders, A., Malapert, P., Alonso, S., François, A., Barrere, C., Seal, R., et al. (2013). TAFA4, a Chemokine-like Protein, Modulates Injury-Induced Mechanical and Chemical Pain Hypersensitivity in Mice. Cell Rep. 5, 378–388.

Dias Carvalho, P., Guimarães, C.F., Cardoso, A.P., Mendonça, S., Costa, Â.M., Oliveira, M.J., and Velho, S. (2018). KRAS Oncogenic Signaling Extends beyond Cancer Cells to Orchestrate the Microenvironment. Cancer Res. 78, 7–14.

Diez del Corral, R., Olivera-Martinez, I., Goriely, A., Gale, E., Maden, M., and Storey, K. (2003). Opposing FGF and retinoid pathways control ventral neural pattern, neuronal differentiation, and segmentation during body axis extension. Neuron 40, 65–79.

Dikovskaya, D., Cole, J.J., Mason, S.M., Nixon, C., Karim, S.A., McGarry, L., Clark, W., Hewitt, R.N., Sammons, M.A., Zhu, J., et al. (2015). Mitotic Stress Is an Integral Part of the Oncogene-Induced Senescence Program that Promotes Multinucleation and Cell Cycle Arrest. Cell Rep. 12, 1483–1496.

Dubrulle, J., McGrew, M.J., and Pourquié, O. (2001). FGF Signaling Controls Somite Boundary Position and Regulates Segmentation Clock Control of Spatiotemporal Hox Gene Activation. Cell 106, 219–232.

Hamburger, V., and Hamilton, H.L. (1992). A series of normal stages in the development of the chick embryo. Dev. Dyn. 195, 231–272.

Henrique, D., Abranches, E., Verrier, L., and Storey, K.G. (2015). Neuromesodermal progenitors and the making of the spinal cord. Development 142, 2864–2875.

Hoshino, R., Chatani, Y., Yamori, T., Tsuruo, T., Oka, H., Yoshida, O., Shimada, Y., Ari-i, S., Wada, H., Fujimoto, J., et al. (1999). Constitutive activation of the 41-/43-kDa mitogen-activated protein kinase signaling pathway in human tumors. Oncogene 18, 813–822.

Kidger, A.M., and Keyse, S.M. (2016). The regulation of oncogenic Ras/ERK signalling by dual-specificity mitogen activated protein kinase phosphatases (MKPs). Semin. Cell Dev. Biol. 50, 125–132.

Kram, D., Henderson, J., Baig, M., Chakraborty, D., Gardner, M., Biswas, S., and Khatua, S. (2018). Embryonal Tumors of the Central Nervous System in Children: The Era of Targeted Therapeutics. Bioengineering 5, 78.

Kuleshov, M.V., Jones, M.R., Rouillard, A.D., Fernandez, N.F., Duan, Q., Wang, Z., Koplev, S., Jenkins, S.L., Jagodnik, K.M., Lachmann, A., et al. (2016). Enrichr: a comprehensive gene set enrichment analysis web server 2016 update. Nucleic Acids Res. 44, W90–97.

Lachmann, A., Torre, D., Keenan, A.B., Jagodnik, K.M., Lee, H.J., Wang, L., Silverstein, M.C., and Ma’ayan, A. (2018). Massive mining of publicly available RNA-seq data from human and mouse. Nat. Commun. 9, 1366.

Le Douarin, N., and Kalcheim, C. (1999). The Neural Crest (Cambridge University Press).

Lunn, J.S., Fishwick, K.J., Halley, P.A., and Storey, K.G. (2007). A spatial and temporal map of FGF/Erk1/2 activity and response repertoires in the early chick embryo. Dev. Biol. 302, 536–552.

Maik-Rachline, G., and Seger, R. (2016). The ERK cascade inhibitors: Towards overcoming resistance. Drug Resist. Updat. 25, 1–12.

Mansour, S.J., Resing, K.A., Candi, J.M., Hermann, A.S., Gloor, J.W., Herskind, K.R., Wartmann, M., Davis, R.J., and Ahn, N.G. (1994). Mitogen-Activated Protein (MAP) Kinase Phosphorylation of MAP Kinase Kinase: Determination of Phosphorylation Sites by Mass Spectrometry and Site-Directed Mutagenesis1. J. Biochem. (Tokyo) 116, 304–314.

Martínez-Morales, P.L., Diez del Corral, R., Olivera-Martínez, I., Quiroga, A.C., Das, R.M., Barbas, J.A., Storey, K.G., and Morales, A.V. (2011). FGF and retinoic acid activity gradients control the timing of neural crest cell emigration in the trunk. J. Cell Biol. 194, 489–503.

Mi, H., Muruganujan, A., Casagrande, J.T., and Thomas, P.D. (2013). Large-scale gene function analysis with the PANTHER classification system. Nat. Protoc. 8, 1551–1566.

Olivera-Martinez, I., Schurch, N., Li, R.A., Song, J., Halley, P.A., Das, R.M., Burt, D.W., Barton, G.J., and Storey, K.G. (2014). Major transcriptome re-organisation and abrupt changes in signalling, cell cycle and chromatin regulation at neural differentiation *in vivo*. Development 141, 3266–3276.

Patel, N.S., Rhinn, M., Semprich, C.I., Halley, P.A., Dollé, P., Bickmore, W.A., and Storey, K.G. (2013). FGF Signalling Regulates Chromatin Organisation during Neural Differentiation via Mechanisms that Can Be Uncoupled from Transcription. PLoS Genet. 9, e1003614.

Reddy, K.B. (2003). [No title found]. Cancer Metastasis Rev. 22, 395–403.

Row, R.H., Tsotras, S.R., Goto, H., and Martin, B.L. (2016). The zebrafish tailbud contains two independent populations of midline progenitor cells that maintain long-term germ layer plasticity and differentiate in response to local signaling cues. Development 143, 244–254.

Samatar, A.A., and Poulikakos, P.I. (2014). Targeting RAS–ERK signalling in cancer: promises and challenges. Nat. Rev. Drug Discov. 13, 928–942.

Shakhova, O. (2014). Neural crest stem cells in melanoma development. Curr. Opin. Oncol. 26, 215–221.

Sneddon, L.U., Halsey, L.G., and Bury, N.R. (2017). Considering aspects of the 3Rs principles within experimental animal biology. J. Exp. Biol. 220, 3007–3016.

Suzuki, A., Makinoshima, H., Wakaguri, H., Esumi, H., Sugano, S., Kohno, T., Tsuchihara, K., and Suzuki, Y. (2014). Aberrant transcriptional regulations in cancers: genome, transcriptome and epigenome analysis of lung adenocarcinoma cell lines. Nucleic Acids Res. 42, 13557–13572.

Tsubota, S., and Kadomatsu, K. (2018). Origin and initiation mechanisms of neuroblastoma. Cell Tissue Res. 372, 211–221.

Wilmerding, A., Rinaldi, L., Caruso, N., Lo Re, L., Bonzom, E., Saurin, A.J., Graba, Y., and Delfini, M.-C. (2021). HoxB genes regulate neuronal delamination in the trunk neural tube by controlling the expression of *Lzts1*. Development 148, dev195404.

Wilson, V., Olivera-Martinez, I., and Storey, K.G. (2009). Stem cells, signals and vertebrate body axis extension. Development 136, 1591–1604.

Won, J.-K., Yang, H.W., Shin, S.-Y., Lee, J.H., Heo, W.D., and Cho, K.-H. (2012). The crossregulation between ERK and PI3K signaling pathways determines the tumoricidal efficacy of MEK inhibitor. J. Mol. Cell Biol. 4, 153–163.

Xie, Z., Bailey, A., Kuleshov, M.V., Clarke, D.J.B., Evangelista, J.E., Jenkins, S.L., Lachmann, A., Wojciechowicz, M.L., Kropiwnicki, E., Jagodnik, K.M., et al. (2021). Gene Set Knowledge Discovery with Enrichr. Curr. Protoc. 1.

